# All in thirty milliseconds: EEG evidence of hierarchical and asymmetric phonological encoding of vowels

**DOI:** 10.1101/482562

**Authors:** Anna Dora Manca, Francesco Di Russo, Francesco Sigona, Mirko Grimaldi

**Affiliations:** Centro di Ricerca Interdisciplinare sul Linguaggio (CRIL), University of Salento, Lecce, Italy; Laboratorio Diffuso di Ricerca interdisciplinare Applicata alla Medicina (DReAM), Lecce, Italy; Dipartimento di Scienze Motorie, Umane e della Salute, University of Rome “Foro Italico”, Rome, Italy; IRCCS Fondazione Santa Lucia, Rome, Italy

**Keywords:** phonological encoding, auditory cortices, electroencephalography, N1, phonemotopic representations

## Abstract

How the brain encodes the speech acoustic signal into phonological representations (distinctive features) is a fundamental question for the neurobiology of language. Whether this process is characterized by tonotopic maps in primary or secondary auditory areas, with bilateral or leftward activity, remains a long-standing challenge. Magnetoencephalographic and ECoG studies have previously failed to show hierarchical and asymmetric hints for speech processing. We employed high-density electroencephalography to map the Salento Italian vowel system onto cortical sources using the N1 auditory evoked component. We found evidence that the N1 is characterized by hierarchical and asymmetric indexes structuring vowels representation. We identified them with two N1 subcomponents: the typical N1 (N1a) peaking at 125-135 ms and localized in the primary auditory cortex bilaterally with a tangential distribution and a late phase of the N1 (N1b) peaking at 145-155 ms and localized in the left superior temporal gyrus with a radial distribution. Notably, we showed that the processing of distinctive feature representations begins early in the primary auditory cortex and carries on in the superior temporal gyrus along lateral-medial, anterior-posterior and inferior-superior gradients. It is the dynamical interface of both auditory cortices and the interaction effects between different distinctive features that generate the categorical representations of vowels.

## 1. Introduction

How does the brain convert the speech acoustic signal into abstract (phonological) representations? According to the *tonotopic principle*, the acoustic structures map directly onto clusters of neurons within the auditory cortex thanks to specific sensitivity of nerve cells to spectral properties of sounds (Romani et al., 1982; Ohl and Scheich, 1997; Saenz and Langers 2014). This seems mostly true for the primary auditory cortex (A1) as well as for the superior temporal gyrus (STG) (Mesgarani et al., 2014). Additionally, the temporal mechanism of auditory encoding, known as the *tonochrony principle*, might augment or supplement the tonotopic strategy in the frequency range critical to human speech: this means that the latency of auditory evoked components is sensitive to some stimulus properties (Roberts et al., 2000).

A classical question is whether speech is processed bilaterally or whether the left hemisphere plays a more dominant role. In line with the *asymmetric sampling in time (*AST) model (Poeppel, 2003), the input speech signal has a bilateral neural representation at the A1 but phonological computations are left lateralized in the ∼20–50 ms temporal integration window while syllabic computation is right lateralized in the ∼150–250 ms integration window in secondary auditory areas. This view is better specified in the *dual-stream model*: at an early stage, a spectro-temporal analysis is carried out in the STG bilaterally. The categorical (phonological) processing, instead, involves the middle to the posterior portion of the superior temporal sulcus (STS) bilaterally, although some indications of left lateralization may emerge (Hickok and Poeppel, 2007; Peelle, 2012). However, the issue remains controversial (Scott and McGettigan, 2013) and the left hemisphere (in particular, the left mid-STG) returned to be dominant for phoneme processing when a meta-analytic investigation of fMRI data was conducted (DeWitt and Rauschecker, 2012).

Further than with other neuroimaging techniques, this issue has been extensively investigated through Magnetoencephalography (MEG) thanks to event-related magnetic fields (ERMFs) and the auditory N1m component (Manca and Grimaldi, 2016). The major challenge is to find a correlation between the hypothesized temporal events (at least three) contributing to the N1/N1m (Näätänen and Picton, 1987; Woods, 1995) and their hierarchical generation from the primary to the secondary auditory areas together with bilateral and/or left hemispheric activation. MEG offers optimal temporal resolution and it is thought that it performs better than Electroencephalography (EEG) in localizing neural activity from the scalp (Baillet, 2017; Ahlfors et al., 2010). Unfortunately, MEG investigations of speech failed to prove both the hierarchical involvement of auditory areas and the clear effects of hemispheric lateralization (Manca and Grimaldi, 2016). When the cortical sources of N1m responses are reported, the supratemporal plane – an area that includes the A1 and the STG (Poeppel et al., 1997; Obleser et al., 2003) – the planum temporale (Obleser et al., 2004a) or the area around the STS (Eulitz et al., 2004) are suggested as the bilateral centers of the speech processing. This limitation might be due to the fact that MEG is particularly sensitive to tangential neuronal sources. Conversely, EEG is sensitive to both radial and tangential sources, although the signal is dominated by radial sources (Malmivuo et al., 1997). Thus, in principle, EEG and the Event-Related Potential (ERP) N1 component should be responsive to a larger range of cortical sources and permit investigators to pick up the dynamical and spatially distributed neuronal activity involved in speech processing. Experiments on the replicability of MEG and EEG measures showed only a minor advantage for MEG (Liu et al., 2002) or even a more EEG accurate localization for the same number of sensors averaged over many source locations and orientations. Furthermore, advances in high-density electrode montages and EEG source analysis have improved the ability to accurately localize EEG signals (Cohen and Halgren, 2003).

We recorded ERPs from 16 subjects and analyzed N1 amplitude, latency, topography and localization of the generators modeled as an equivalent current dipole (ECD) in searching for the neural indexes of the auditory cortex generating vowels representations. We investigated the five-vowel system characterizing the Salento Italian (SI) variety spoken in Southern Apulia: /i, ε, a, ɔ, u/. This simple phonological system is suitable for our aim since it results the most common vowel system in the world’s languages (de Boer, 2001). Thus, the findings of relative N1 modulations, due to the peculiar structures of this vowel system, may provide evidence concerning the universal neural mechanisms for vowel representations.

According to linguistic theory (Halle, 2002; Stevens, 2002), the relevant representational linguistic primitives are not vowels and consonants, but rather smaller units: i.e., *distinctive features*. Distinctive features are universal representational links between articulatory plans and acoustic outputs and must have correlates in terms of both articulation and audition. Bundles of distinctive features, characterized by polar oppositions (binary values), form the consonant and vowel segments. For instance, vowels features identify binary contrasts for tongue height and backness/frontness in the mouth or lip rounding. So, distinctive features specify the phonemic contrasts that are used in the language, such that a change in the value of a feature can contrastively generate a new word: e.g., English /i:/ [+high] in [’bi:d] *bead* vs. /ε/ [-high, -low] in [’bεd] *bed*. The auditory pathways decode the speech signal structures and ensure the identification of acoustic landmarks providing evidence for the action of specific articulators and contrastive features marking phonemes (**Fig. 1**). Vowels are characterized by the first two peaks of their spectral envelopes (F1 and F2 values in Hz): F1 inversely correlates with the tongue height (low F1 are consistent with high vowels), while F2 correlates with tongue frontness in the mouth (high F2 values are consistent with front vowels) and lip rounding (lip rounding lowers the F2 values) (Stevens, 2002; Peterson & Barney, 1952). Thus, the five-vowel system under investigation is marked by three contrasts for Height (referring to the vertical tongue position in the mouth): [+high] /i/, /u/; [-high, -low] /ε/, /ɔ/; [+low] /a/; and one contrast for Place (referring to the horizontal tongue position in the mouth): [-back] /i/, /ε/; [+back] /a/, /ɔ/, /u/. The [±round] feature is redundant since /ɔ/, /u/ are both [+back] and [+round] and the vowel /a/ is contrastively only [+low], so its features, [+back, -round], are predictable by means of this specification (Calabrese, 1995) (**Fig. 1** and **Table 1**).

**Table 1.** Pitch (F0), Formant Frequency (F1, F2, F3 in Hz) mean values and rise and fall-times (ms) of the vowels used as stimuli (standard deviation is given in parenthesis). The parameters F2-F1 are also given.

| Vow. | F0 | F1 | F2 | F3 | F2-F1 | Rise | Fall |
| --- | --- | --- | --- | --- | --- | --- | --- |
| [i] | 145 | 294 (±11) | 2325 (±50) | 2764 (±28) | 2031 (±47) | 21 | 28 |
| [ε] | 145 | 549 (±32) | 1880 (±46) | 2489 (±60) | 1330 (±71) | 40 | 32 |
| [a] | 140 | 794 (±30) | 1231 (±24) | 2528 (±95) | 418 (±46) | 33 | 27 |
| [ɔ] | 140 | 550 (±14) | 856 (±13) | 2551 (±54) | 306 (±21) | 29 | 22 |
| [u] | 130 | 310 (±12) | 660 (±33) | 2437 (±49) | 349 (±25) | 22 | 23 |

**Fig. 1.**
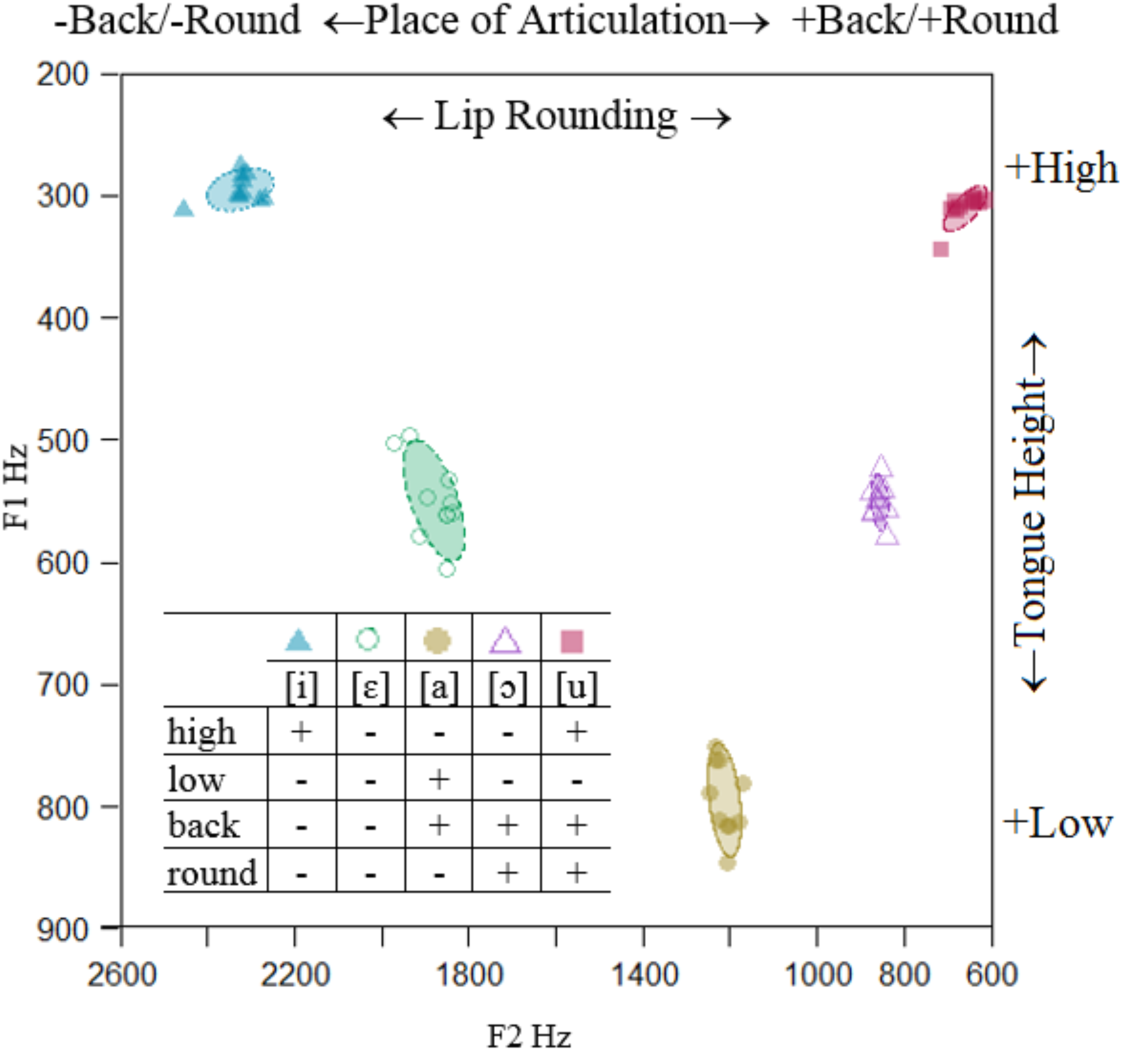
68.27% confidence ellipse corresponding to ±1 SD from the bivariate mean. The F1 is inversely correlated with articulatory tongue height, while the F2 reflects the place of articulation in the horizontal dimension.

Whether speech sound maps are solely determined by bottom–up acoustic information or whether they are modulated by top–down information relying on abstract featural information is another question left open by linguistic research and MEG studies (Manca & Grimaldi, 2016). A solid evidence is that the acoustic distance between the first two formants of a vowel is preserved in the auditory cortex and is directly reliable in sensor and source data along the Talairach 3D coordinate system: lateral medial (*x*), anterior-posterior (*y*), and inferior-superior (*z*) gradients. At the same time, amplitudes, latencies and spatial gradients in the auditory cortices tentatively suggest that acoustic-articulatory properties are affected by top-down features such as Height, Place and Round (Manca & Grimaldi, 2016). Clues of orderly cortical representations of abstract features emerge when more than one couple of vowels are investigated (Obleser et al., 2004a; Ohl and Scheich, 2004), or when an entire phonological system has been studied with appropriate statistical analyses able to discern different levels of auditory brain operations (Scharinger et al., 2011). Thus, it is hard to disambiguate between N1m evidences suggesting pure acoustic patterns and those indicating abstract phonological features.

With the aim to assess the hierarchical involvement of auditory areas, the effects of hemispheric lateralization, and the acoustic/abstract featural representations of SI vowels, we investigated the N1 ERP component employing two linear mixed effects statistical models: the acoustic model included the predictors F1 and F2 as fixed effects and Subject as random intercept; the phonological model included the phonological predictors Height (three contrasts) and Place (one contrast) as fixed effects and Subject as random intercept. In this way, we want to ascertain whether spatial arrangement of neuronal sources are a pure bottom-up reflection of spectro-temporal differences between vowels or whether they are simultaneously warped by top-down information relying on polar oppositions determined by abstract distinctive features information.

## 2. Materials and Methods

### 2.1 Subjects

Sixteen volunteer students of the University of Salento (8 females; mean±SD, 23±3 years) participated in the experiment after providing written informed consent. All subjects were consistently right-handed according to Handedness Edinburgh Questionnaire (Oldfield, 1971) and none of them had any known neurological disorders or other significant health problems. The Ethical Committee of the *Vito Fazzi Hospital* in Lecce approved the experimental procedure. The study was carried out in accordance with the guidelines of the <u>Declaration of Helsinki</u>. The data were acquired in the *Centro di Ricerca Interdisciplinare sul Linguaggio (*CRIL) in Lecce (Italy).

### 2.2 Stimuli and procedure

The stimuli consisted of the five stressed SI vowels and a pure tone. A native Italian male speaker (age 32) realized ten repetitions of each vowel in isolation, at a normal rate. The speech signal was recorded in a soundproof room with CSL 4500 and a Shure SM58-LCE microphone with a sampling rate of 44.1 kHz and an amplitude resolution of 16 bits. The stimuli were edited and analyzed by using the speech analysis software Praat 5.2 (Boersma and Weenink, 2011). All stimuli were normalized for duration (200 ms), for the F0 values according to the values of a representative sample of SI vowels (Grimaldi, 2009): i.e., 130Hz for /i/ 140Hz for /ε, a, ɔ/, and 145Hz for /u/, and for intensity (70 dB/SPL). The F0-F3 formant values were measured in the vowel steady tract (0.025 s) centered at the midpoint. The ramp for rise/fall times was not edited to preserve the natural sounding speech (**Table 3**) as it has been showed that the rise- and fall-times times of vowels do not affect the relative N1 latencies and amplitudes (Grimaldi et al., 2016; Gage et al., 1998). A pure tone of 1000 Hz and duration of 200 ms was created by Praat software. In the experimental protocol the best five exemplars of each vowel type and the pure tone were binaurally transmitted to the subjects through two loudspeakers (Creative SBS 2.1 350) at a comfortable loudness (about 70dB/SPL) with Presentation software 2.0. Before the EEG recordings, participants familiarized with the stimuli. All the subjects were able to identify each of the vowels with an accuracy of 100%.

### 2.3 Experimental Design

During the experiment, the participants were seated in front of a computer monitor in a shielded room. They were asked to listen to the vowels and to push a button with their left index finger whenever they heard a pure tone of 1 KHz used as distractor stimulus (**Fig. 2)**. Two blocks of 1000 vowel stimuli were each presented. Each block consisted of 200 tokens per vowel category and 70 distractor stimuli. Stimuli were randomly presented with a variable inter-stimulus interval that ranged between 1000 ms to 1400 msec. The distractor stimulus was interspersed with a probability between 6% and 7% in the train of the vowel sounds. To reduce excessive eye movements, participants were asked to fixate on a white crosshair located in the center of the monitor. The experiment lasted approximately 1hour.

**Fig. 2.**
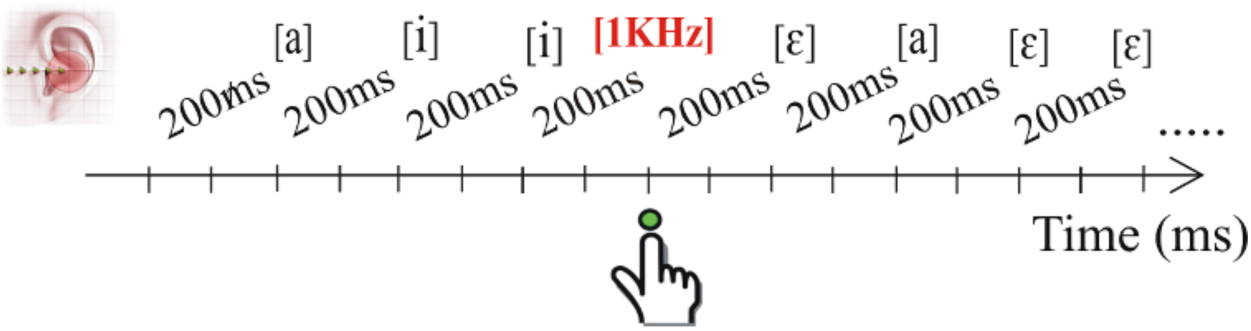
Scheme of the experimental design. Participants had to press a response button when they heard a pure tone (occurring with a probability between 6% and 7%), represented as [1Khz].

### 2.4 Data acquisition and preprocessing

Continuous EEG was recorded by using a 64-channel ActiCap^TM^ (Brain Products GmbH, Germany) and Brain Vision Recorder 1.20 (Brain Products GmbH, Germany) at a sampling rate of 250 Hz, an online band pass filter of 0.16-80 Hz, and a notch filter at 50 Hz. Vertical eye movements were monitored using Fp2 and an additional electrode attached below the right eye. FT9 and FT10 were used for horizontal movements. The online reference was at FCz, the ground was AFz, and the impedance was kept under 5 KΩ.

Off-line signal processing was carried out using Brain Vision Analyzer 2.0.1 (Brain Products GmbH, Germany). The EEG was segmented in relation to the onset of the five vowels: thus, the distractor and the following stimulus were left out of analyses. ERP epochs of 1200 ms (including 200 ms pre-stimulus baseline) were extracted, digitally filtered by a 1–30 Hz band pass filter (48db) and re-referenced to the average of the left and right mastoids (M1/2). Ocular artifacts were removed by applying an ICA algorithm and, additionally, rejection criteria for trials were set to 120 µV maximum absolute difference. Artifact-free segments were separately averaged for each vowel and a baseline correction was applied over the applied pre-stimulus portion. Finally, grand averages were computed across all subjects and for each vowel type. Analyses were focused on the N1 component. The grand-average scalp distribution revealed two N1 distinct peaks in the range between 80 to 160 ms here termed N1a and N1b for convenience. The earliest N1a component was identified as the most prominent peak between 125 and 135 ms after the stimulus onset on central medial electrodes. The later N1b peak was identified as the most prominent peak between 145 and 155 ms over left central, fronto-central and centro-parietal electrodes.

### 2.4 Source analysis

Tridimensional topographical maps and an estimation of the N1a and N1b intracranial sources were conducted using BESA 2000. We used the spatiotemporal source analysis of BESA that estimates location, orientation and time course of the equivalent dipolar sources (ECD) by calculating the scalp distribution obtained for a given model (forward solution). This distribution was then compared to that of the actual AEPs. Interactive changes in source location and orientation lead to the minimization of residual variance between the model and the observed spatiotemporal AEP distribution. The three-dimensional coordinates of each ECD in the BESA model were determined with respect to the Talairach axes. BESA assumed a realistic approximation of the head (based on the MRI template based on 24 subjects). The possibility of interacting ECDs was reduced by selecting solutions with relatively low ECD moments with the aid of an “energy” constraint (weighted 20% in the compound cost function, as opposed to 80% for the residual variance). The optimal set of parameters was found in an iterative manner by searching for a minimum in the compound cost function. Latency ranges for fitting (N1a: 100-125 ms; N1b: 125-155 ms) were chosen to minimize overlap between the two, topographically distinctive components. A first model was made on the grand-average AEP to obtain a reliable and stable model of the N1a and N1b for all vowels using two bilateral mirror symmetric pairs of ECDs on the basis of the topographical maps obtained here and in previous studies (McDonald et al., 2003; Teder-Sälejärvi et al., 2002; Teder-Sälejärvi et al. 2005) showing bilateral distribution on the auditory N1 component. Then, to compare statistically the N1 source localizations across vowels, the model was used as starting point to model the AEP of each subject fitting the source locations and orientation on the individual data. The accuracy of the source model was evaluated by measuring its residual variance as a percentage of the signal variance, as described by the model, and by applying residual orthogonality tests (ROT) (Bocker et al., 1994). The resulting individual time series for the ECD moments (the source waves) were subjected to an orthogonality test, referred to as a source wave orthogonality-test (SOT) (52). All t-statistics were evaluated for significance at the 5% level.

### 2.5 Statistical analysis

The N1 peak amplitudes and latencies were measured at the most prominent electrodes: e.g., Cz or CPz for the N1a and at C3 or CP3 for the N1b. The latency and amplitude values were analyzed separately for each N1 component with two linear mixed effects model using R (R Core Team, 2015), *lme4 (*Bates et al., 2015), and *multcomp (*Hothorn et al., 2008) (with Tukey post-hoc). The acoustic model included the acoustic predictors F1 and F2 as fixed effects and Subjects as random intercept; the phonological model included the phonological predictors Height and Place as fixed effects and Subjects as random intercept. Specifically, we defined three contrasts for Height ([+high] /i, u/, [-high, -low] /ε, ɔ/, and [+low] /a/), one for Place ([-back] /i, ε/, [+back] /u, ɔ, a/). We separated spectro-temporal and phonological predictors in two different models as, notwithstanding the correlation existing between formants and distinctive features, we wanted to determine whether the N1 amplitudes, latencies and ECD sources are better accounted for by acoustic gradient predictors or by distinctive features predictors.

Furthermore, we tested the hemispheric asymmetries for the N1b on the mean amplitudes for the four strongest electrodes in each hemisphere: i.e., C3-FC5-F3-FC1 for the left and C4-FC6-F4-FC2 for the right hemisphere. The acoustic and phonological models were built by using Hemisphere and the Acoustic (F1 and F2) or the Height and Place predictors as fixed effects and Subjects as random effect. Visual inspection of residual plots did not reveal any evidence of deviations from homoscedasticity or normality. P values were obtained by likelihood ratio tests of the full model with effect in question against the model without that effect. We performed model comparison analysis on the base of previous literature (Baayen, 2008; Pinheiro and Bates, 2000) to investigate what model exhibits the best fit for the data. The best model will be the one with lower values of the Akaike Information Criterion (AIC) and Bayes Information Criterion (BIC), while the statistical significance (α=0.05) was evaluated using likelihood ratios (which provided as logarithm units, logLR) that are associated with a p-value.

## 3. Results

### 3.1 Waveforms and topographical maps

In the N1 range, two distinct negative components were detected named here N1a and N1b for convenience. The N1a peaked at 125-135 ms on medial electrodes around the vertex, while the N1b at 145-155 ms over central, fronto-central and centro-parietal electrodes of the left hemisphere. **Fig. 3** shows AEP waveforms of representative electrodes where the activity was prominent and **Fig. 4** the topographical mapping of the two components elicited by each vowel. The N1a topography coincided with the classical N1 distribution focusing on medial centro-parietal scalp areas with a tangential distribution (negative on the vertex and positive on bilateral temporal sites). This component was posterior for /a/ and /ɔ/ to /ε/, /i/, /u/. The N1b was observed over the left parieto-temporal scalp with a radial distribution at the skull (its positive counterpart was not detectable from the scalp). The N1b was more lateral and posterior for the [+low] /a/ and the [-high -low] /ε, ɔ/ to the [+high] /i, u/.

**Fig. 3.**
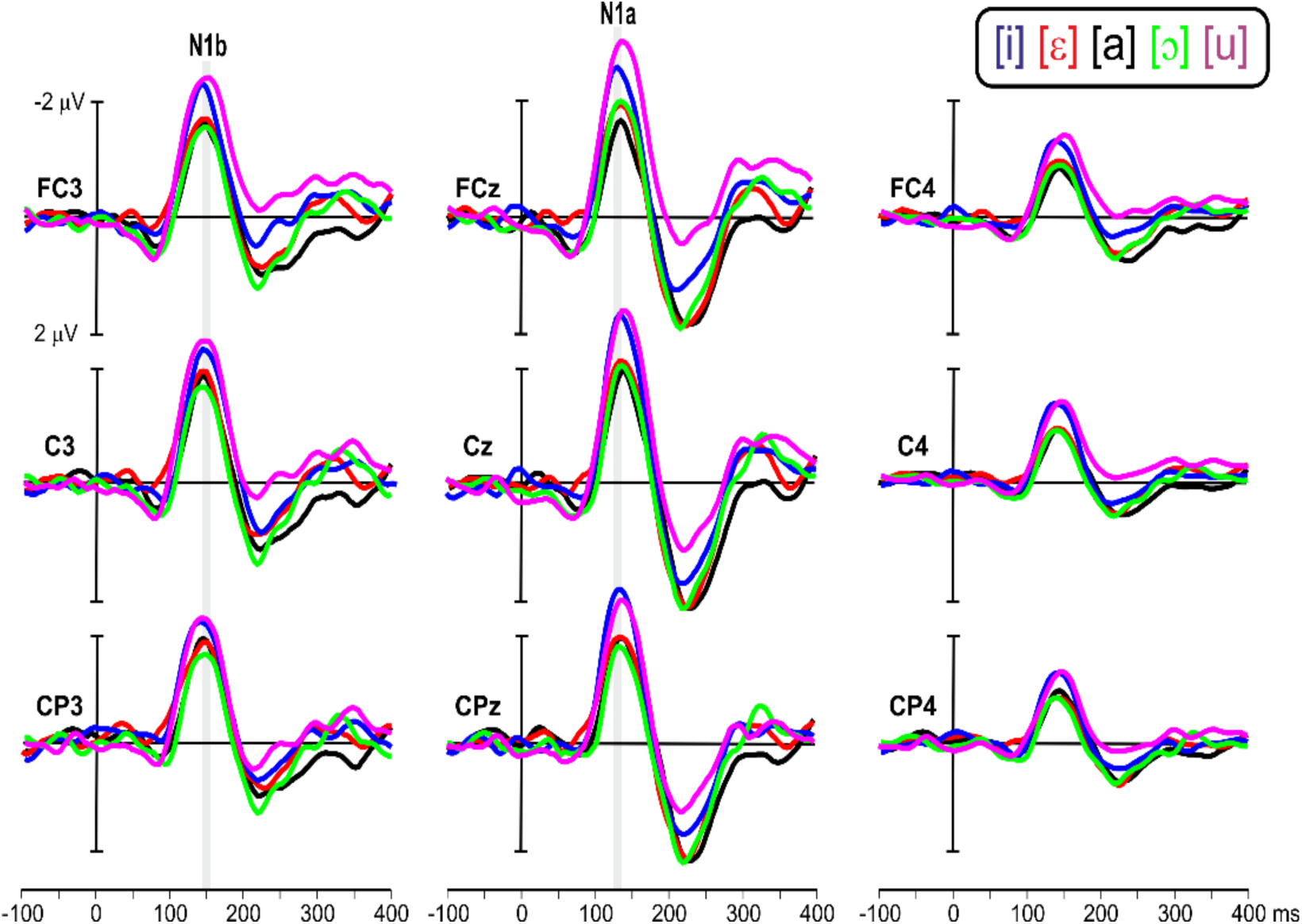
Grand average (N=11) of the N1a and N1b components at the most representative electrode sites. The five vowels are superimposed using different colors. The time windows in which the two N1 subcomponents peak are marked by a vertical gray bar.

**Fig. 4.**
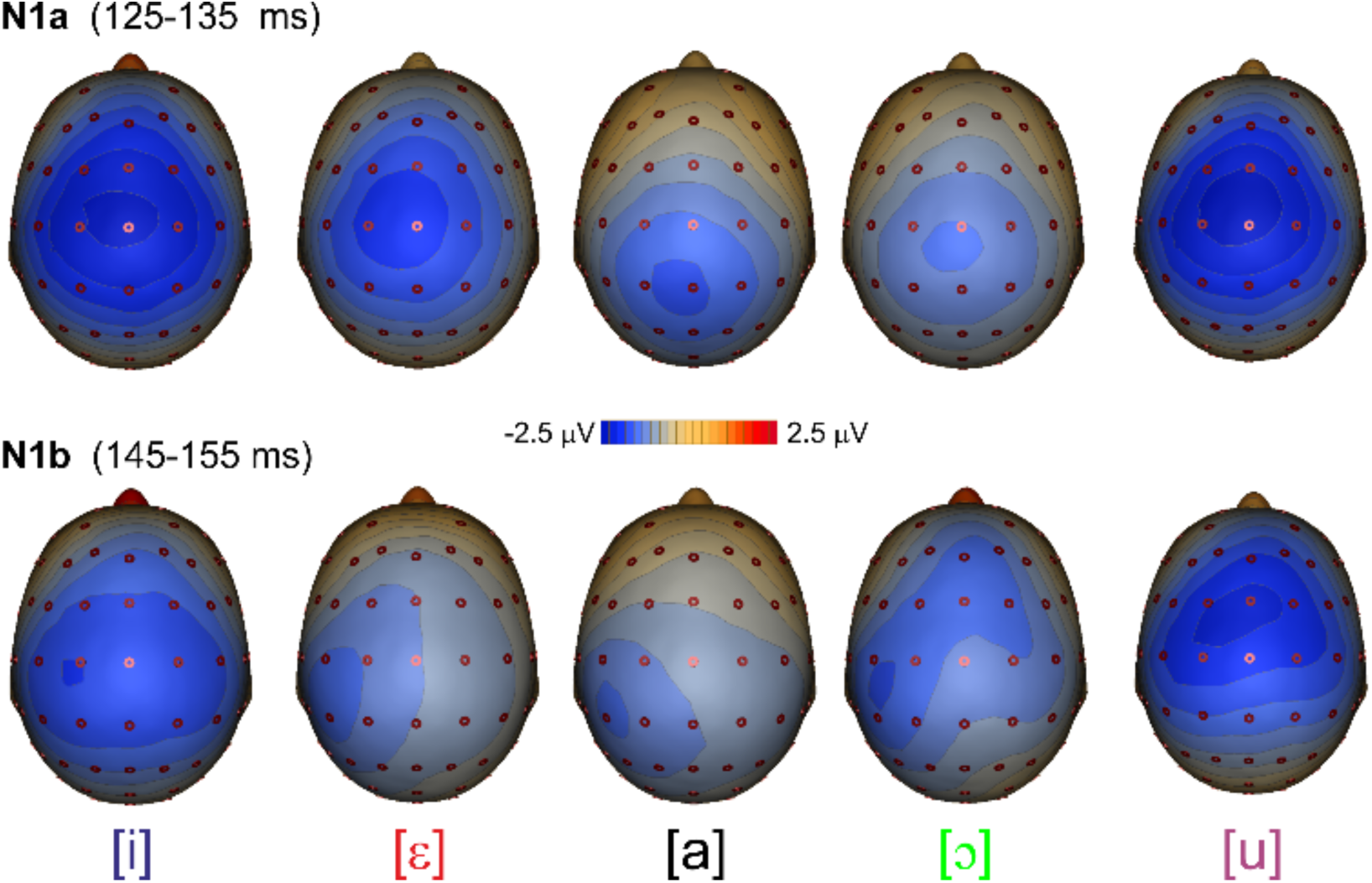
N1a and N1b topographical three-dimensional maps displayed from a top view.

### 3.2 Hemispheric asymmetry

To test hemispheric asymmetries on the N1b amplitudes, the Hemisphere effect was added in the models. Both acoustic (χ^2^(1) = 91.1, P < 0.001) and phonological (χ^2^(1) = 93.9, P < 0.001) models showed a leftward laterality. On average, N1b amplitudes were -2.32 µV (SD±0.5) for the left and -1.11 µV (SD±0.3) for the right. The main effects of F1 (χ^2^(1) = 4.6, P = 0.031) and Height (χ^2^(2) =8.9, P = 0.011) and the interactions Hemisphere x F1 (χ^2^(1) = 5.8, P = 0.016) and Hemisphere x Height (χ^2^(2) = 6.0, P = 0.048) were statistically relevant. In the left hemisphere the [+high] /i, u/ vowels, with low F1, elicited greater responses than the [-high -low] /ε, ɔ/ and [+low] /a/ vowels (P < 0.001). F2 (χ^2^(1) = 21.5, P = 0.643) and Place (χ^2^(1) = 96.1, P = 0.327) were not statistically relevant. Model comparison revealed that the phonological model provides a better fit for the data to the acoustic model (logLR = 2.444).

### 3.3 Amplitudes and latencies

The N1a and N1b amplitude and latency values are shown in **Table 2** and displayed in **Fig. 5 (A, B)**. In **Fig. 5C** is represented the leftward laterality of N1b in respect of the N1a. For both N1a and N1b amplitudes, the acoustic model showed a main effect for F1 (N1a: χ^2^(1) = 10.5, P = 0.001; N1b: χ^2^(1) = 7.4, P =0.006) and the phonological model a main effect for Height (N1a: χ^2^(2) = 1.2,1 P = 0.; N1b: χ^2^(2) = 8.3, P = 0.015). That is, the amplitudes increase with decreasing F1 values of vowels generating contrasts for Height (cfr. **Fig. 1**).

**Table 2.**
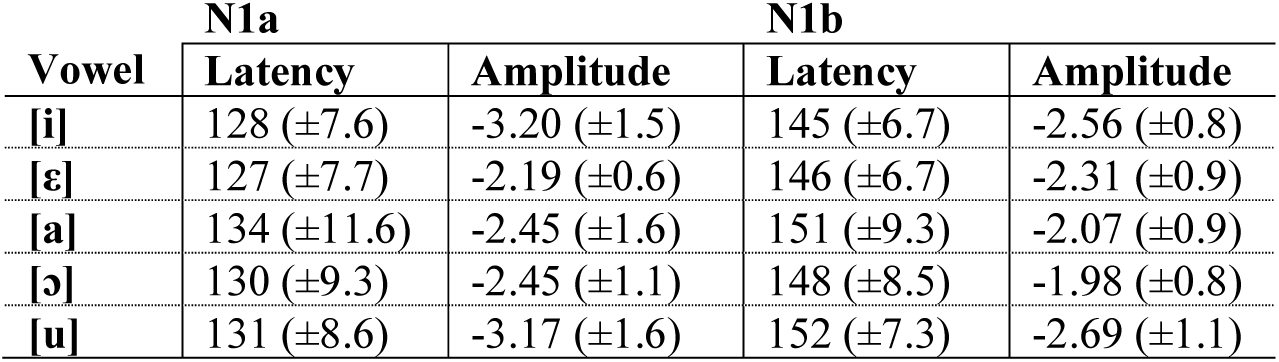
Mean amplitude (μV), latency (ms), and standard deviation (±) values of the five SI vowels for the N1a and N1b for N=11.

**Fig. 5.**
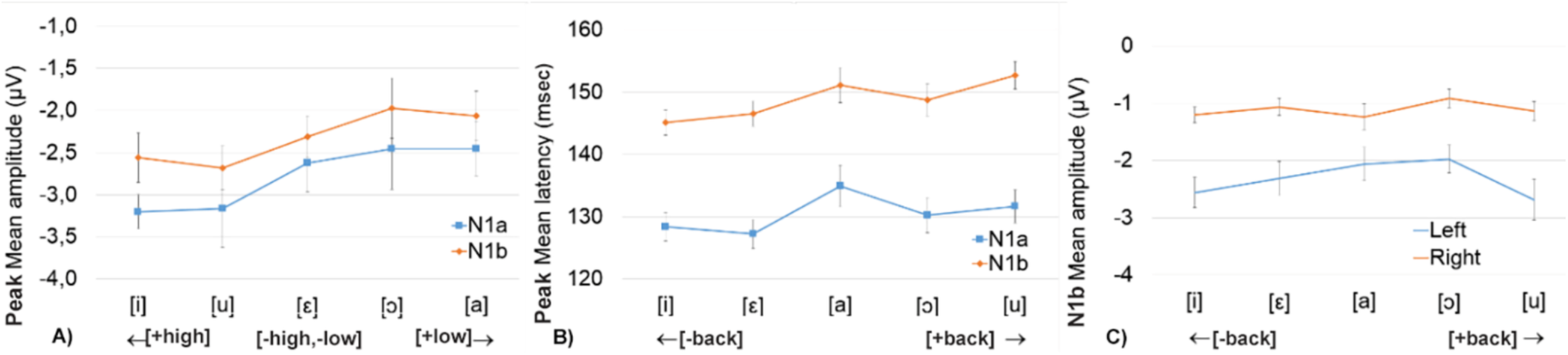
Amplitudes, latencies, and hemispheric asymmetries of the N1 components. A) mean amplitudes of the two N1 subcomponents for [+high] /i/, /u/, [-high, -low] /ε/, /ɔ/ and [+low] /a/ vowels; B) mean latency of the two N1 subcomponents for [+back] /a, ɔ, u/ and [-back] /i, ε/, vowels; C) hemispheric asymmetries in the amplitude of the N1b component as function of the perceived vowels.

In the phonological model, the N1a responses to the [+high] /i, u/ elicited greater amplitudes than the [-high, -low] /ε, ɔ/ (P = 0.003) and [+low] /a/ (P = 0.020); however, the /ε, ɔ, a/ vowels did not statically differ (P > 0.001). These findings were partially paralleled by the N1b: responses to /i, u/ elicited greater amplitude than /ε, ɔ/, but responses to /i, u/ were not different from /a/ responses (P=0.081); again, the vowels [ε], [ɔ] and [a] did not statically differ (P > 0.992). F2 and Place were not statistically relevant (N1a: F2 (χ^2^(1) = 966, P = 0.756; Place (χ^2^(2) = 52.9, P = 0.467; N1b: F2 (χ^2^(1) = 5090, P = 0.943; Place (χ^2^(2) = 3.7, P = 0.540). The phonological model provided a better fit for the N1a data (logLR = 1.176) whereas the acoustic model provided a better fit for the N1b data (logLR = 0.837).

As for latency, the acoustic model did not show significant effects for the N1a data (F1: (χ^2^(1) = 3.1, P = 0.077; F2: (χ^2^(1) = 3.5, P = 0.059). The phonological model bared a better goodness of fit (logLR = 3.708) showing a significant effect for Place (χ^2^(1) = 4.8, P = 0.028): the [+back] /a, ɔ, u/ were, on average, 3.12 ms later than the [-back] /i, ε/. Height was not statistically relevant (χ^2^(1) = 5.9, P = 0.050). Statistics for the N1b values showed a main effect for F2 (χ^2^(1) = 9.0, P = 0.003) and Place (χ^2^(1) = 7.7, P = 0.005) confirming that the [+back] vowels with low F2 values were later to the [-back] vowels (on average 4.9 ms). The F1 and Height predictors were not statistically relevant (F1: (χ^2^(1) = 0.0413, P = 0.839; Height: (χ^2^(2) = 0.59.7, P = 0.742). The acoustic model fitted slight better than phonological model (logLR= 0.151).

### 3.4 ECD localization

**Table 3** shows the source coordinates for the five vowels and the two components. The intracranial localization of the N1 sources for the five vowels are shown in **Fig. 6**. The waves represent the time course of those sources in both hemispheres (averaged across vowels). For all vowels, the N1a was bilaterally localized within primary auditory cortex in the Brodmann area (BA41). The N1a time-course showed that this component initiated at 80 ms and peaked at 130 ms with equal intensity in the two hemispheres. The N1b was localized more ventrally and anteriorly within the STG in the BA22. The N1b time-course revealed that this component initiated at 110 ms and peaked at 150 ms and that was much larger in the left hemisphere than in the right (t(10)=23.4, P < 0.0001).

**Table 2.** Talairach coordinates of the bilateral source locations of the five vowels (and relative mean) for the N1a and N1b components. The ± symbols before the X values indicate that sources were constrained to be symmetric in both hemispheres.

| Vowel | N1a |  |  | N1b |  |  |
| --- | --- | --- | --- | --- | --- | --- |
|  | X | Y | Z | X | Y | Z |
| [i] | ±46 | -24 | 13 | ±52 | -4 | 1 |
| [ε] | ±48 | -26 | 12 | ±49 | -8 | -2 |
| [a] | ±54 | -30 | 15 | ±60 | -25 | -2 |
| [ɔ] | ±51 | -22 | 9 | ±56 | -17 | -7 |
| [u] | ±50 | -17 | 10 | ±46 | -11 | -10 |
| Mean | ±50 | -24 | 12 | ±53 | -13 | -4 |

**Fig. 6.**
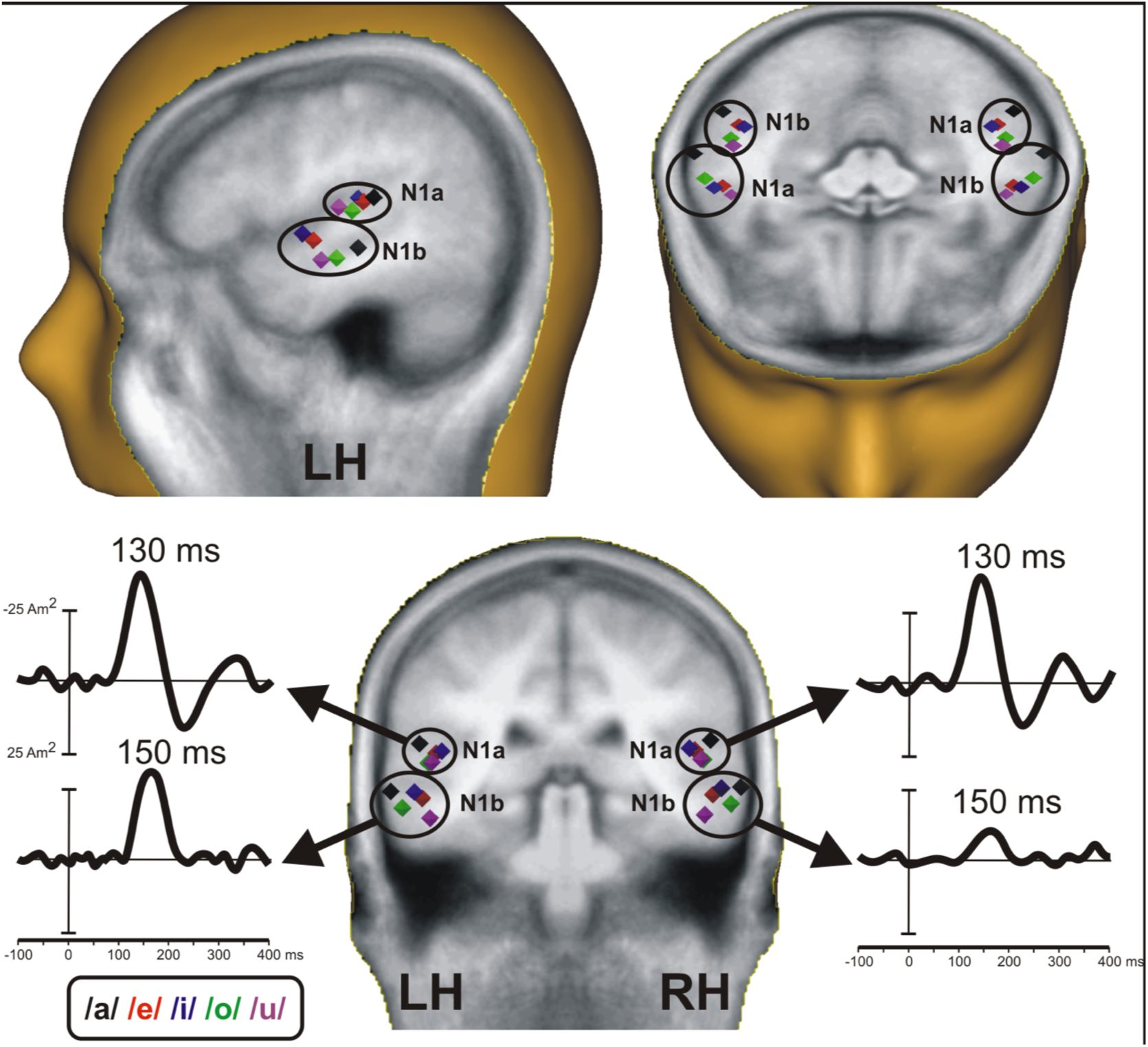
Source locations and time-course of the N1a and the N1b. Vowel representation is coded by different colours (blue, red, black, green and purple dots respectively). LH=left hemisphere, RH=right hemisphere.

In **Fig. 7** the N1a and N1b bivariate source distribution along the XY and ZY axes are represented in two-dimensional planes.

#### 3.4.1 Lateral-medial dimension (x)

The acoustic model for the N1a ECD showed F1 (χ^2^(1) = 9.82, P = 0.002) and F2 (χ^2^(1) = 6.49, P = 0.011) effects: this suggest that the /a, ɔ, u/ vowels close in the F2-F1 dimension (i.e. [+back] vowels]) elicited lateral source locations to the /ε, i/ vowels with larger inter-format distances (i.e. [-back] vowels). In the phonological model effects for Height (χ^2^(2) = 7.07, P = 0.029) and Place (χ^2^(1) = 8.40, P = 0.004) emerged. The Height effect indicated that the sources of the [-low] /a/ was on average 4 mm lateral to the [+high] /i, u/ (P = 0.022), whereas the other vowels were not cortically distinguished (P > 0.001); the Place effect evidenced that the [+back] vowels were more lateral than the [-back] vowels. The phonological model fit better for the N1a data (logLR = 1.619). For the N1b data, the acoustic model showed a main effect for F1 (χ^2^(1) = 31.5, P < 0.001) and F2 (χ^2^(1) = 1.26, P = 0.262): highest F1 values (e.g. /a/) were collocated anteriorly and lowest F2 values (e.g. /u/) medially. The phonological model evidenced effects for Height (χ^2^(2) = 27, P < 0.001). Post-hoc showed that the [+low] /a/ was at the most lateral position – on average 7 mm to [-high, -low] /ε, ɔ/ and 12 mm to [+high] /i, u/ – and that the [-high, -low] vowels were 4 mm lateral to the [+high] vowels (P < 0.001). The interaction Height × Place (χ^2^(1) = 29.2 P < 0.001) showed that: (i) /a/ and /ɔ/ were not cortically distinguished; (ii) within the [-high -low] vowels, /ε/ was medial to /ɔ/; (iii) within the [+high] vowels, /i/ was lateral to /u/. The phonological model fits better the N1b data (logLR = 14.5).

#### 3.4.2 Anterior-posterior dimension (y)

The N1a acoustic model showed effects for F1 (χ^2^(1) =76.1, P < 0.001) and F2 (χ^2^(1) =51.7, P < 0.001): they indicated that vowels with high F1 (i.e. /a/) and vowels with high F2 (i.e. /ε/ and /i/) tended to elicit ECDs at posterior locations in respect of /ɔ/ and /u/. In the phonological model a main effect for Height (χ^2^(2) = 84.8, P < 0.001) and Place (χ^2^(1) = 45.9, P < 0.001) emerged. On average, the [+low] /a/ was at the most posterior position – on average 8 mm to the [-high, -low] /ε, ɔ/ and 11 mm to the [+high] /i, u/ (P < 0.001); in their turn, the [-high, -low] /ε, ɔ/ were 3 mm posterior to the [+high] /i, u/. Moreover, the [+back] vowels /ɔ, u/ were anterior to the [-back] vowels /ε, i/. The interaction Height × Place (χ^2^(1) = 4.2, P= 0.038) revealed Place effects within the [-high, -low] and [+high] vowels: so that /ε/ was posterior to /ɔ/ and /i/ was posterior to /u/. The phonological model better described the source data (logLR=2.86).

With regard to the N1b data, the acoustic model revealed effects for the F1 (χ^2^(1) = 96.4, P < 0.001) and F2 (χ^2^(1) =47.8, p < 0.001): this means that vowels close in the F2-F1 dimension (i.e., the [+back] /a, ɔ, u/) elicited posterior ECDs to vowels with larger inter-formant distances (i.e., [-back] /ε, i/). In the phonological model, we found that Height (χ^2^(2) = 111, P <0.001) and Place (χ^2^(1) = 89.2, P < 0.001) predictors affected the ECD patterns. On average, the [+low] /a/ was at the most posterior location – about 8 mm to [-high, -low] /ε, ɔ/ and 13 mm to [-high] /i, u/; in their turn, /ε, ɔ/ were 3 mm posterior to /i, u/ (P <0.001). Crucially, contrary to the N1a sources, the [+back] /a, ɔ, u/ were on average 8 mm posterior to the [-back] /ε, i/. In addition, the interaction of Height × Place (χ^2^(1) = 5.3, p = 0.021) indicated the Place effects within [-high, -low] and [+high] vowels: contrary the N1a sources, the [-back] /ε/ was anterior to the [+back] /ɔ/ and the [-back] /i/ was anterior to the [+back] /u/. The phonological model fitted the data better than the acoustic model (logLR = 22.0).

#### 3.4.3 Inferior-superior dimension (z)

The N1a was affected by F1 (χ^2^(1) =18.5, P < 0.001) and F2 (χ^2^(1) = 23.9, P < 0.001) in the acoustic model: vowels with higher F1 and higher F2 values (e.g., /a/, /ε/, and /i/) tended to be generated in the superior ECDs. The phonological model provided a better fit for the data (logLR = 16.13) evidencing a main effect for Height (χ^2^(2) = 59.9, P < 0.001): the [+low] /a/ was located at the most superior location (P < 0.001), but the [-high, -low] /ε, ɔ/ were not cortically distinguished from the [+high] /i, u/ (P = 0.111). Also, an effect for Place was evident: the [+back] /a, ɔ, u/ were on average 3 mm inferior to the [-back] /i, ε/ (χ^2^(1) = 37.1, P < 0.001).

Moving to the N1b, the acoustic model highlighted clear effects for F1 (χ2(1) = 25.7, P <0.001) and F2 (χ2(1) = 96.2, P < 0.001): again, this suggest that the vowels /a, ɔ, u/, with F2-F1 close dimension, were inferior to the vowels /i, ε/ with larger F2-F1 distances. In the phonological model, the Place effect was significant (χ2(1) = 79.1, P < 0.001): the [+back] vowels were inferior to the [-back] vowels by 8 mm on average. Moreover, a significant interaction Height x Place was noticeable (χ2(1) = 22.4, P < 0.001): within the [+back] vowels, [ɔ] is superior to [u]; within the [- back] vowels, [ε] is inferior to [i]. The vowels [a] and [ε] were not cortically separated (P = 0.92), while [a] was statistically different from [ɔ]. The phonological model fitted better the data (logLR =3.47).

## 4. Discussion

Three are the novel findings of the present study. First of all, we found evidence for different hierarchical indexes structuring vowel representation within the N1 component: the N1a peaking at 125-135 ms in the A1 (BA41) with a tangential distribution and the N1b peaking at 145-155 ms in the STG (BA22) with a radial distribution. Secondly, these components are characterized by hemispheric asymmetries: the N1a shows a bilateral activity while the N1b shows a leftward activity. Finally, the hierarchical and hemispheric modulation of the N1a-N1b shed light on the encoding of spectro-temporal properties of vowels into distinctive features representations through the tonotopic activation of lateral-medial, anterior-posterior, and inferior-superior gradients.

### 4.1 The origins of the N1a and N1b components

According to literature, the scalp distribution of the N1 responses to clicks, noise, bursts, and tones hint at least three distinct subcomponents (Näätänen and Picton, 1987). The first subcomponent is maximally recorded from the fronto-central scalp, peaks between 85-110 ms and is generated by tangentially orientated currents in both A1 (Hari et al. 1980; Wood and Wolpaw, 1982); the second subcomponent is detectable approximately at 150 ms in the mid temporal scalp regions and is generated by radially oriented neuronal sources in the STG, and the third subcomponent is a negative wave at the vertex at 100 ms whose generators are not known (Wood and Wolpaw, 1982; Wolpaw and Penry, 1975; Picton et al., 1078; Inui et al., 2006). The N1a and N1b we found present patterns that are in accordance with the first and the second subcomponent previously hypothesized (according to the Wood’s list (Woods, 1995), the N1’/P90 and N1c, respectively). However, our N1a shows a slight longer peak latency than previous data. This may depend on the nature of our stimuli: actually, vowels have complex spectro-temporal properties whose recognition might require more cognitive efforts than non-speech stimuli. As far as we know, this is the first study that found clear evidence for these hypothesized subcomponents in humans and, importantly, for speech sounds. Probably, early MEG studies failed to report the different intracranial origin on the N1 subcomponents as the N1m sources were generally modelled by a single ECD in each hemisphere. Conversely, we used two bilateral mirror symmetric pairs of ECDs on the basis of the topographical maps obtained, which permitted us to capture the multiple (temporally-differentiated) N1 cortical generators. Also, it is very likely that the EEG sensitivity to radial and tangential ECDs as compared to MEG, which is blind to radially oriented ECDs (Eulitz et al., 2004; Malmivuo et al., 1997), permitted us to separate the activity related to vowel encoding. To deeply understand the hemispheric modulation of these components we need to discuss amplitudes, latency and especially source data.

### 4.2 The acoustic and phonological models: amplitudes, latency and sources data

As in ways observed before, amplitudes show broad F1 and Height encoding processes in both N1a and N1b: amplitudes increase with decreasing F1 values so that the [+high] vowels /i, u/ elicited greater amplitudes than non-high vowels (Obleser et al., 2003; Scharinger et al., 2011; Shestakova et al., 2004). For latencies, the acoustic model revealed a significant effect only for the N1b: vowels with low F2 (/a, ɔ, u/) were later to vowels with high F2 (/i, ε/). Instead, the phonological model show effects for both N1a and N1b revealing that the [+back] /a, ɔ, u/ peaked later than the [-back] /i, ε/ (Obleser et al., 2004a,b). Crucially, the phonological model fit better the N1a while the acoustic model fit better the N1b amplitudes and latencies suggesting that Height and Place distinctive features are early encoded in the A1.

Source data offer a fine-grained picture. Previous studies showed that N1m ECDs are dependent on both spectro-temporal cues and distinctive features (Manca and Grimaldi, 2016). The lateral-medial axis showed medial locations for sounds with high frequencies or lateral positions for close F2-F1 distances, so that the [+back] vowels (with small F2-F1 interval) result more lateral than the [-back] vowels (Eulitz et al., 2004; Obleser et al., 2004b); also, the [+round] vowels (with low F2) elicit more lateral sources (Scharinger et al., 2011). The anterior-posterior plane seems responsive to F1 and F2 values associated with Height and Place: so, the [+high] vowels result more anterior than the [-high] vowels and the [+back] vowels more posterior than the [-back] vowels (Obleser et al., 2004a; Scharinger et al., 2011). The inferior-superior axis showed sensitivity to F1 and Height: it has been found that low vowels are superior to high vowels (Obleser et al., 2003) but also the reverse pattern seems true only for the [-back] vowels (Scharinger et al., 2012). Yet, the sources of rounded vowels turn out to be inferior to non-round vowels (Scharinger et al., 2011). We replicated these tonotopic data adding new patterns thanks to the N1a-N1b hierarchical-hemispheric modulation.

Of note is the fact that the phonological model provides a better fit both for N1a and N1b ECDs along all tonotopic gradients. This suggests that: (i) distinctive features are better predictors than acoustic F1 and F2 patterns for vowel representations; (ii) representation processes begin early in the A1 in both hemispheres (Bernal and Ardila, 2016) and carry on in the left STG. The acoustic model provides a general result concerning the encoding of the F2-F1 relation that specifies vowels for Place: tongue retraction lowers F2 frequencies reducing F2-F1 distances. So, as in early studies (Eulitz et al., 2004; Obleser et al., 2004b), the [+back] vowels /a, ɔ, u/, with close F2-F1, result more lateral in the N1a, more posterior and more inferior in the N1b than the [-back] vowels /i, ε/ (**Fig. 7**). This finding is also in line with an ECoG study that investigated the phonological American English system (Mesgarani et al., 2014): the STG electrodes showed a selective response to F2-F1 differences separating low-back, low-front and high-front vowels. In our study, the ECD patterns found for the F2-F1 parameters are preserved within the phonological model which shows that the [+back] vowels are more lateral in the N1a, more posterior in the N1b, and more inferior in the N1b. However, both for our and previous studies the acoustic model fails to adequately capture the tonotopic mapping of the three tongue heights marking the SI vowel system (as well as the American English vowel system which differentiates also between tense and lax vowels). Our phonological model caught these contrasts and more importantly elucidated their nature thanks to Height x Place interactions affecting source ECDs (interactions never found even when the same statistical models here adopted were employed with MEG (Scharinger et al., 2011) to study the Turkish vowel system).

**Fig. 7.**
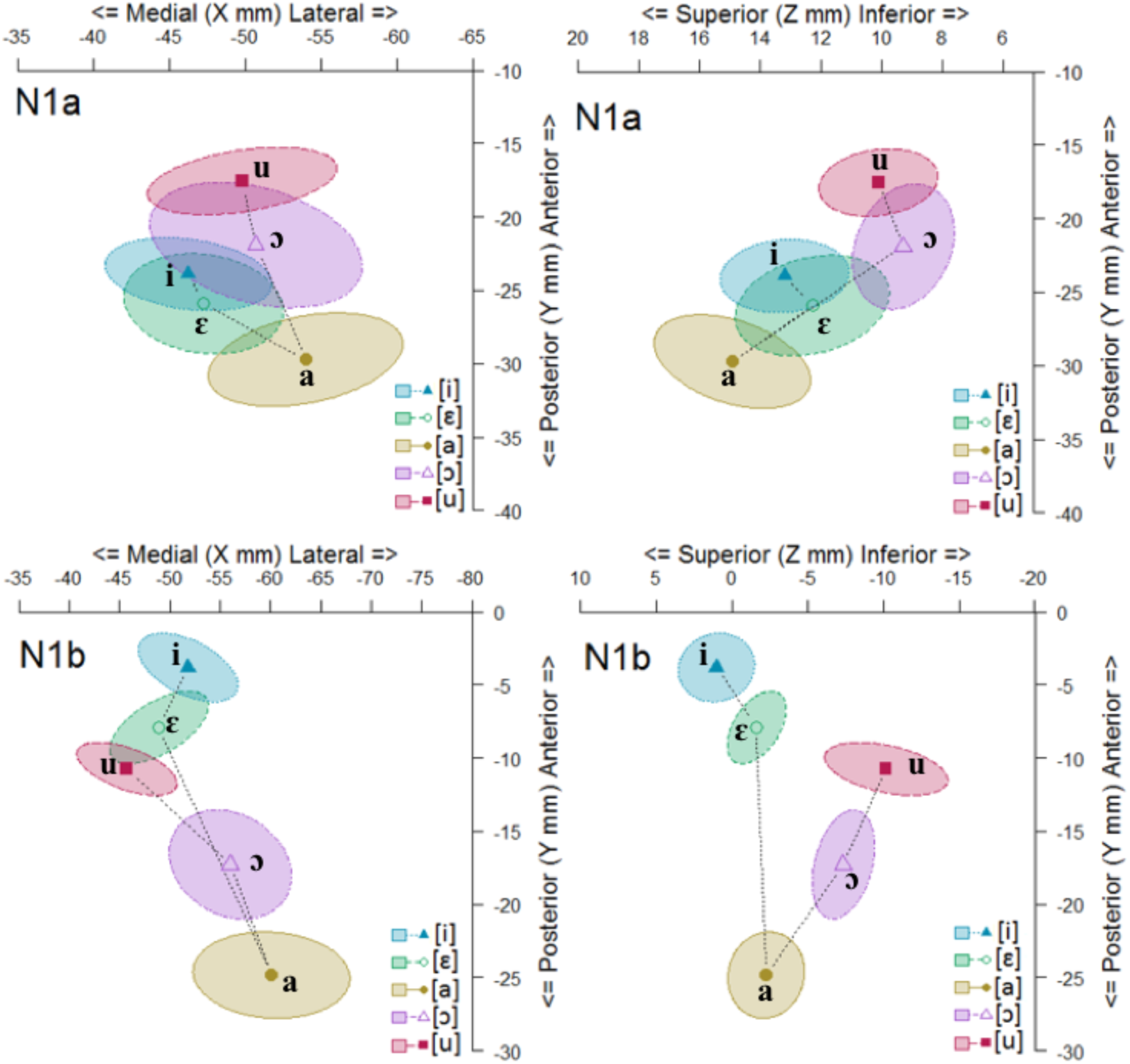
N1a (upper part) and N1b (lowest part) source locations in the two-dimensional plane determined by the Medial-Lateral/Posterior-Anterior (on the left) and Superior-Inferior/Posterior-Anterior axes (on the right). Mean values are represented by dots. 68.27% confidence ellipses of the source location for each vowel (corresponding to ±1 SD from the bivariate mean).

The N1a and N1b anterior-posterior gradients highlight that the [+low] /a/ results at the most posterior position and the [-high, -low] /ε, ɔ/ is significantly posterior to the [+high] /i, u/. Crucially, the interaction Height x Place effects reveal source ECDs selectivity within the [-high, -low] and [+high] vowels: in the N1a, the [-back] /ε/ is posterior to the [+back] /ɔ/ and the [-back] /i/ is posterior to the [+back] /u/. In the N1b the reverse pattern holds because the /ε, i/ vowels reach a more anterior position during the N1a-N1b hierarchical-hemispheric modulation (**Fig. 7**): /ε/ is anterior to /ɔ/, and /i/ is anterior to /u/ (as already found for the Turkish system (Scharinger et al., 2011). A further modulation Height x Place is noticeable in the inferior-superior gradient where again, moving from the N1a to the N1b, the /ε, i/ vowels reach a superior position (**Fig. 7**). This modulation selectively separate ECDs for Height contrasts within the [+back] /ɔ, u/ and [-back] /ε, i/: the [-high, -low] /ɔ/ is superior to the [+high] /u/ and the [-high, -low] /ε/ is inferior to the [+high]/i/.

Overall, our results suggest that the computational processes leading to abstract representations of SI vowels warp the spatial arrangement of neuronal sources according to distinctive features so that: (i) Place features are encoded along the lateral-medial (N1a), anterior-posterior (N1b), and inferior-superior (N1b) gradients separating [±back] vowels; (ii) the three contrasts for Height are encoded along the N1a-N1b anterior-posterior gradient; (iii) within this gradient the [-high, -low] /ε, ɔ/ and [+high] /i, u/ are additionally separated for Place through N1a-N1b hierarchical-hemispheric modulations; (iv) finally, the same vowels are also separated for Height along the N1b inferior-superior gradient. Briefly, for SI vowels it seems that at first a general mapping for Place and Height features is performed; then, further encoding for Height and Place are executed to specify [±back] features within the [-high, -low] vowels and the [±high] features within [-back] and [+back] vowels separately.

### 4.3 Vowels encoding and hemispheric lateralization

Our findings have important implications for current theories on speech hemispheric lateralization. The data above discussed contrast with models hypothesizing that spectro-temporal analysis of speech sounds is bilaterally performed in the A1 (Poeppel et al., 2006) and then phonological computations are left lateralized, but also with models suggesting that the spectro-temporal analysis is carried out in the STG bilaterally while phonological processing in the left STS (Hickok and Poeppel, 2007) or with other point of views stressing the exclusive contribution of the left STG in phonological representations (Scott and McGettigan, 2013; DeWitt and Rauschecker, 2012; Scott et al., 2000; Scott and Johnsrude, 2003). We maintain that the initial stage of speech encoding is bilaterally performed in the A1; however, we showed that at this level the cortical spatial arrangement is already warped by phonemotopic patterns according to distinctive features principles. So, it is probable that spectro-temporal analysis, previously attributed to the A1, is peculiar to the cochlea-brainstem pathways (until the proximity of the A1): here properties of the speech waveform are mirrored with remarkable fidelity (Bidelman et al., 2013). Conversely, late cortical evoked activities, from the A1 to the STG, progressively encodes the phonological features necessary to generate categorical speech percepts. Indeed, our data suggest that the bilateral A1 and the left STG forms multiple (parallel) representations of vowels leading to the conversion of the acoustic signal into categorical patterns through hierarchical reshaping of neuronal maps along the lateral-medial, anterior-posterior, and inferior-superior gradients: this dynamical interface between the A1 and the STG generates the encoding of Place and Height features for SI vowels.

## 5. Conclusion

Overall, the findings of the present study suggest that vowel discretization is the result of a continuous process that converts physical states into other physical states (Grimaldi, in press). We hypothesize that the spectro-temporal states characterizing vowels are continuously converted into appropriate neurophysiological states: in this way, properties of the spectro-temporal states undergo changes interacting with the neurophysiological states until synchronized synapses, phonemotopically distributed within the A1 and the STG, are generated. From this perspective, the classical distinction between bottom-up processes reflecting acoustic differences and top-down processes reflecting distinctive features representations should be reinterpreted as a continuous-dynamical process involving changes of physical states (spectro-temporal states into neurophysiological states) where progressive structure and property rearrangements result in categorical representation of vowels according to distinctive features specifications.

## Author Contributions

M.G. conceived the study and leaded the research. A.D.M., M.G. and F.D.R. designed the experiment. A.D.M. acquired the data. F.D.R. and A.D.M. analyzed the EEG data. F.S. developed and executed the statistical analysis. M.G., A.D.M, F.D.R. and F.S. interpreted the results. M.G., A.D.M., F.D.R. and F.S. wrote the manuscript.

## Acknowledgements

this research was supported by the Italian Ministry of Education, University and Research, Grant. No. 20128YAFKB_006.

## Data availability

The data that support the findings of this study are available on request from the corresponding author M.G. The data are not publicly available due to a prohibition by the Italian law. Juridical restrictions set by the Italian law prevent public access to the collected data, be it anonymized or non-anonymized, when data are recorded from human individuals. As the consent given by the subjects only applies to the specific study reported in our manuscript, no portion of the data collected could be used or released for use by third parties.

---

**Figure 2**: on the top: grand average (N=11) of the N1a and N1b components at the most representative electrode sites. The five vowels are superimposed using different colors. The time windows in which the two N1 subcomponents peak are marked by a vertical gray bar. On the bottom: N1a and N1b topographical three-dimensional maps displayed from a top view.

**Figure 3**: A) mean amplitudes of the two N1 subcomponents for [+high] /i/, /u/, [-high, -low] /ε/, /ɔ/ and [+low] /a/ vowels; B) mean latency of the two N1 subcomponents for [+back] /a, ɔ, u/ and [-back] /i, ε/, vowels; C) hemispheric asymmetries in the amplitude of the N1b component as function of the perceived vowels.

**Figure 4**: on the top: source locations and time-course of the N1a and the N1b. Vowel representation is coded by different colours (blue, red, black, green and purple dots respectively). LH=left hemisphere, RH=right hemisphere. On the bottom: N1a (upper part) and N1b (lowest part) source locations in the two- dimensional plane determined by the Medial-Lateral/Posterior-Anterior (on the left) and Superior- Inferior/Posterior-Anterior axes (on the right). Mean values are represented by dots. 68.27% confidence ellipses of the source location for each vowel (corresponding to ±1 SD from the bivariate mean).

## References

Ahlfors, S. P., Han, J., Belliveau, J. W., Hämäläinen, M. S. 2010. Sensitivity of MEG and EEG to Source Orientation. Brain. Top. 23, 227–232.

Baayen, R. H. 2008. Analyzing Linguistic Data. A practical introduction to statistics, Cambridge University Press, Cambridge.

Baillet, S. 2017. Magnetoencephalography for brain electrophysiology and imaging. Nat. Neur. 20, 327–339.

Bates, D., Maechler, M, Bolker, B. 2015. Walker S Fitting Linear Mixed-Effects Models Using lme4. J. Stat. Soft. 67(1): 1–48. DOI:10.18637/jss.v067.i01.

Bernal, B., Ardila, A. 2016. From Hearing Sounds to Recognizing Phonemes: Primary Auditory Cortex is A Truly Perceptual Language Area. AIMS Neur. 3(4), 454–473. DOI:10.3934/Neuroscience.2016.4.454.

Bidelman, G. M., Moreno, S., Alain, C. 2013. Tracing the emergence of categorical speech perception in the human auditory system. NeuroIm. 79, 201–212.

Bocker, K.B.E, Cornelis, H.M., Brunia, C.H.M., Van den Berg-Lenss, M.M.C. 1994. A Spatiotemporal dipole model of the stimulus preceding negativity prior to feedback stimuli. Br. Top. 7, 71–88.

Boersma, P., Weenink, D. 2011. Praat: doing phonetics by computer (Computer program), Version 5.2. http://www.praat.org/.

Calabrese, A. 1995. Constraint-based theory of Phonological markedness and simplification procedures. Ling. Inqu. 2(26), 373–463.

Cohen, D., Halgren, E. 2003. Magnetoencephalography (Neuromagnetism): in Encyclopedia of Neuroscience. Elsevier, Amsterdam, pp. 615–622.

de Boer, B. 2001. The Origins of Vowel Systems. Oxford University Press, Oxford.

DeWitt, I., Rauschecker, J.P. 2012 Phoneme and word recognition in the auditory ventral stream. Proc. Natl. Acad. Sci. USA 109(8), 505–514.

Diesch, E, Eulitz, C., Hampson, S., Ross, B. 1996. The neurotopography of vowels as mirrored by evoked magnetic field measurements Bra. & Lang. 53, 143–168.

Eulitz, C., Obleser, J., Lahiri, A. 2004, Intra-subject replication of brain magnetic activity during the processing of speech sounds. Cog. Br. Res. 19, 82–91.

Gage, N., Poeppel, D., Roberts, T.P.L., Hickok, G. 1998. Auditory evoked M100 reflects onset acoustics of speech sounds. Br. Res. 814(1), 236–239.

Grimaldi, M. 2009. Acoustic correlates of phonological microvariations. The case of unsuspected highly diversified metaphonetic processes in a small area of Southern Salento (Apulia): in Tock, D., Wetzels, W. L., (Eds.),Romance Languages and Linguistic Theory 2006, Benjamins, Amsterdam, pp. 89–109.

Grimaldi, M. in press. The phonetics-phonology relationship in the neurobiology of language: in Petrosino, P, Cerrone, P. van der Hulst, H., (Eds.), From sounds to structures: beyond the veil of Maya, De Gruyter, Berlin, pp. 66–104.

Grimaldi, M., Manca, A.D., Di Russo, F. 2016. Electroencephalography evidence of vowels computation and representation in auditory cortex: in Di Sciullo, A. M. (Ed.), Biolinguistic Investigations on the Language Faculty, Benjamins, Amsterdam, pp 80–100.

Halle, M. 2002. From memory to speech and back: Papers on phonetics and phonology 1954–2002. Mouton de Gruyter, Berlin.

Hari, R., Aittoniemi, K., Järvinen, M. L., Katila, T., Varpula, T. 1980. Auditory evoked transient and sustained magnetic fields of the human brain localization of neural generators. Ex. Brain Res. 40(2), 237–240.

Hickok, G., Poeppel, D. 2007. The cortical organization of speech processing. Nat. Neur. 8, 393–402.

Hothorn, T., Bretz, F., Westfall, P. 2008. Simultaneous Inference in General Parametric Models. Biom. J. 50(3), 346–363.

Inui, K., Okamoto, H., Miki, K., Gunji, A., Kakigi, R. 2006. Serial and parallel processing in the human auditory cortex: A magnetoencephalographic study. Cer. Cor. 16, 18–30.

Liu, A. K., Dale, A. M., Belliveau, J. W. 2002. Monte Carlo simulation studies of EEG and MEG localization accuracy. Hum. Brain Map. 16, 47–62.

Malmivuo, J., Suihko, V., Eskola, H. 1997. Sensitivity distributions of EEG and MEG measurements. Biom. Eng. IEEE Trans. 44, 196–208.

Manca, A. D., Grimaldi, M. 2016. Vowels and Consonants in the Brain: Evidence from Magnetoencephalographic Studies on the N1m in Normal-Hearing Listeners. Front. Psychol. 7,1413. DOI:10.3389/fpsyg.2016.01413.

McDonald, J.J., Teder-Sälejärvi, W.A., Di Russo, F., Hillyard, S.A. 2003. Neural substrates of perceptual enhancement by crossmodal spatial attention. J. Cog. Neur. 15, 10–19.

Mesgarani, N., Cheung. C., Johnson, K., Chang, E. F. 2014. Phonetic feature encoding in human superior temporal gyrus. Science 343, 1006–1010.

Näätänen, R., Picton, T. 1987. The N1 wave of the human electric and magnetic response to sound: A review and an analysis of the component structure. Psychoph. 24, 375–425.

Obleser, J., Elbert, T., Eulitz, C. 2004b. Attentional influences on functional mapping of speech sounds in human auditory cortex. BMC neuroscience 5, 24. https://doi.org/10.1186/1471-2202-5-24.

Obleser, J., Elbert, T., Lahiri, A., Eulitz, C. 2003. Cortical representation of vowels reflects acoustic dissimilarity determined by formant frequencies. Cog. Bra. Res. 15, 207–213.

Obleser, J., Lahiri, A., Eulitz, C. 2004a. Magnetic brain response mirrors extraction of phonological features from spoken vowels. J. Cog. Neur. 16, 31–39 DOI:10.1162/089892904322755539.

Ohl, F. W., Scheich, H. 1997. Orderly cortical representation of vowels based on formant interaction. Proc. Natl. Acad. Sci. USA 94, 9440–9444.

Oldfield, R,C, 1971. The assessment and analysis of handedness: the Edinburgh inventory. Neuropsycho. 9(1): 97–113.

Peelle, J. E. 2012. The hemispheric lateralization of speech processing depends on what “speech” is: A hierarchical perspective. Front. Hum. Neur. 6, 309. DOI:10.3389/fnhum.2012.00309.

Peterson, G. E., Barney, H. L. 1952. Control methods used in a study of the vowels. J. Acou. Soc. Am. 242, 175–184.

Picton, T. W., Campbell, K. B., Baribeau-Braun, J., Proulx, G. B. 1978. The neurophysiology of human attention: A tutorial review: in Requin, J. (Ed.), Attention and performance VII, Erlbaum, Hillsdale, New Jersey, pp. 429–467.

Pinheiro, J., Bates, D.M. 2000. Linear and nonlinear mixed-effects models in S and S-Plus. Springer-Verlag, New York.

Poeppel, D. 2003. The analysis of speech in different temporal integration windows: Cerebral lateralization as ‘asymmetric sampling in time’. Speech. Comm. 41, 245–255.

Poeppel, D. et al. 1997. Processing of vowels in supratemporal auditory cortex. Neur. Lett. 221, 145–148 DOI:10.1016/S0304-3940(97)13325-0.

R Core Team, 2015. R: A language and environment for statistical computing. R Foundation for Statistical Computing, Vienna, Austria. http://www.R-project.org/.

Roberts, T. P., Ferrari, P., Stufflebeam, S.M., Poeppel, D. 2000. Latency of the auditory evoked neuromagnetic field components: Stimulus dependence and insights toward perception. Jour. Clin. Neur. 17, 114–129.

Romani, G. L., Williamson, S. J., Kaufman, L. 1982. Tonotopic organization of the human auditory cortex. Science 216, 1339–1340.

Saenz, M., Langers, D.R.M. 2014. Tonotopic mapping of human auditory cortex. Hear. Res. 307, 42–52.

Scharinger, M., Idsardi, W. J., Poe, S. A. 2011. Comprehensive Three-dimensional Cortical Map of Vowel Space. J. Cog. Neur. 23, 3972–3982.

Scharinger, M., Monahan, P. J., Idsardi, W. J. 2012. Asymmetries in the processing of vowel height. J. Spec. Lang. Hear. Res. 55(3): 903–918.

Scott, S. K., Blank, C. C., Rosen, S., Wise, R. J. S. 2000. Identification of a pathway for intelligible speech in the left temporal lobe. Bra. J. of Neur. 123(12), 2400–2406 http://dx.doi.org/10.1093/brain/123.12.2400.

Scott, S. K., Johnsrude, I. S. 2003. The neuroanatomical and functional organization of speech perception. Tren. Neur. 26, 100–107.

Scott, S.K., McGettigan, C. 2013. Do temporal processes underlie left hemisphere dominance in speech perception? Bra. & Lang 127, 36–45.

Shestakova, A., Brattico, E., Soloviev, A., Klucharev, V., Huotilainen, M. 2004. Orderly cortical representation of vowel categories presented by multiple exemplars. Br. Cog. Res. 21, 342–350.

Stevens, K. N. 2002. Toward a model for lexical access based on acoustic landmarks and distinctive features. J. Acou. Soc. Am. 111, 1872–1891.

Teder-Sälejärvi, W.A., McDonald, J.J., Di Russo, F., Hillyard, S.A. 2002. An analysis of audio-visual crossmodal integration by means of event related potential ERP) recordings. Cog. Br. Res. 14, 106–114.

Teder–Sälejärvi, W.A., Di Russo, F. McDonald, J.J., Hillyard, S.A. 2005. Effects of Spatial Congruity on Audio-Visual Multimodal Integration. J. Cog. Neur. 17, 1396–1409.

Wolpaw, J.R., Penry, J. K. A. 1975. Temporal component of the auditory evoked response. Electr. Clin. Neuroph. 39, 609–620.

Wood, C. C., Wolpaw, J. R. 1982. Scalp distribution of human auditory evoked potentials. II. Evidence for multiple sources and involvement of auditory cortex. Electr. Clin. Neuroph. 54, 25–38.

Woods, D. L. 1995. The component structure of the N 1 wave of the human auditory evoked potential. Electr. Clin. Neuroph-Supp. 44, 102–109.].

